# sNucDrop-Seq: Dissecting cell-type composition and neuronal activity state in mammalian brains by massively parallel single-nucleus RNA-Seq

**DOI:** 10.1101/154476

**Authors:** Peng Hu, Emily Fabyanic, Zhaolan Zhou, Hao Wu

## Abstract

Massively parallel single-cell RNA sequencing can precisely resolve cellular diversity in a high-throughput manner at low cost, but unbiased isolation of intact single cells from complex tissues, such as adult mammalian brains, is challenging. Here, we integrate sucrose-gradient assisted nuclear purification with droplet microfluidics to develop a highly scalable single-nucleus RNA-Seq approach (sNucDrop-Seq), which is free of enzymatic dissociation and nucleus sorting. By profiling ~11,000 nuclei isolated from adult mouse cerebral cortex, we demonstrate that sNucDrop-Seq not only accurately reveals neuronal and non-neuronal subtype composition with high sensitivity, but also enables analysis of long non-coding RNAs and transient states such as neuronal activity-dependent transcription at single-cell resolution *in vivo*.

---

A fundamental challenge in deciphering cellular composition and cells’ functional states in complex mammalian tissues manifests in the extraordinary diversity of cell morphology, size and local microenvironment. While current high-throughput single-cell RNA-Seq approaches have proved to be powerful tools for interrogating cell types, dynamic states and functional processes *in vivo* ^1^, these methods require the preparation of intact, single-cell suspensions from freshly isolated tissues, which is only practical for easily-dissociated embryonic and young postnatal tissues. This requirement poses an even greater challenge for cells with complex morphology such as mature neurons. Harsh enzymatic treatment not only favors recovery of easily dissociated cell types, but also introduces aberrant transcriptional changes during the dissociation process ^2^. In addition, skeletal and cardiac muscle cells are frequently multinucleated and are large in size. For instance, each adult mouse skeletal muscle cell contains hundreds of nuclei and is ~5,000 μm in length and 10-50 μm in width ^3^. Thus, existing high-throughput single-cell capture and library preparation methods, including isolation of cells by fluorescence activated cell sorting (FACS) into multi-well plates, sub-nanoliter wells, or droplet microfluidic encapsulation, are not optimized to accommodate these unusually large cells. Isolating individual nuclei for transcriptome analysis is a promising strategy, as single-nucleus RNA-Seq methods avoid strong biases against cells of complex morphology and large size ^2, 4-6^, and can be potentially standardized to accommodate the study of various tissues. However, current single-nucleus RNA-Seq methods rely on fluorescence-activated nuclei sorting (FANS) ^4, 5^ or Fluidigm C1 ^6^ to capture nuclei, and thus cannot easily be scaled up to generate a comprehensive atlas of cell types in a given tissue, much less a whole organism.

An ideal solution to increase the throughput of single-nucleus RNA-Seq is to integrate nucleus purification with massively parallel single-cell RNA-Seq methods such as Drop-Seq ^7^, InDrop ^8^, or equivalent commercial platforms (e.g. 10x Genomics ^9^). However, single-nucleus RNA-Seq is currently not supported on these droplet microfluidics platforms. Inhibitory effects due to cellular debris contamination and/or inefficient lysis of nuclear membranes might contribute to this failure. Historically, nuclei of high purity can be isolated from solid tissues or from cell lines with fragile nuclei by centrifugation through a dense sucrose cushion to protect nucleus integrity and strip away cytoplasmic contaminants. The sucrose gradient ultracentrifugation approach has been adapted to isolate neuronal nuclei for profiling histone modifications ^10^, nuclear RNA ^11^, and DNA methylation ^11, 12^ at genome-scale. To test whether this nuclei purification method supports single-nucleus RNA-Seq analysis, we isolated nuclei from cultured cells, as well as freshly isolated or frozen adult mouse brain tissues through douncing homogenization followed by sucrose gradient ultracentrifugation (**Fig. 1a and Supplementary Fig. 1**). After quality assessment and nuclei counting, we performed emulsion droplet barcoding of the nuclei and library preparation with both Drop-Seq and 10x Genomics platforms. While the10x Genomics single-cell 3’ solution workflow supports cDNA amplification only from whole cells (possibly due to inefficient lysis of nuclear membrane), the Drop-Seq platform yielded high quality cDNA and sequencing libraries from both whole cells and nuclei (freshly isolated or frozen samples) (**Supplementary Fig. 2**). These results suggest that nucleus purification and nuclear membrane lysis are critical factors for efficient library preparation in single-nucleus RNA-Seq.

**Figure 1.**
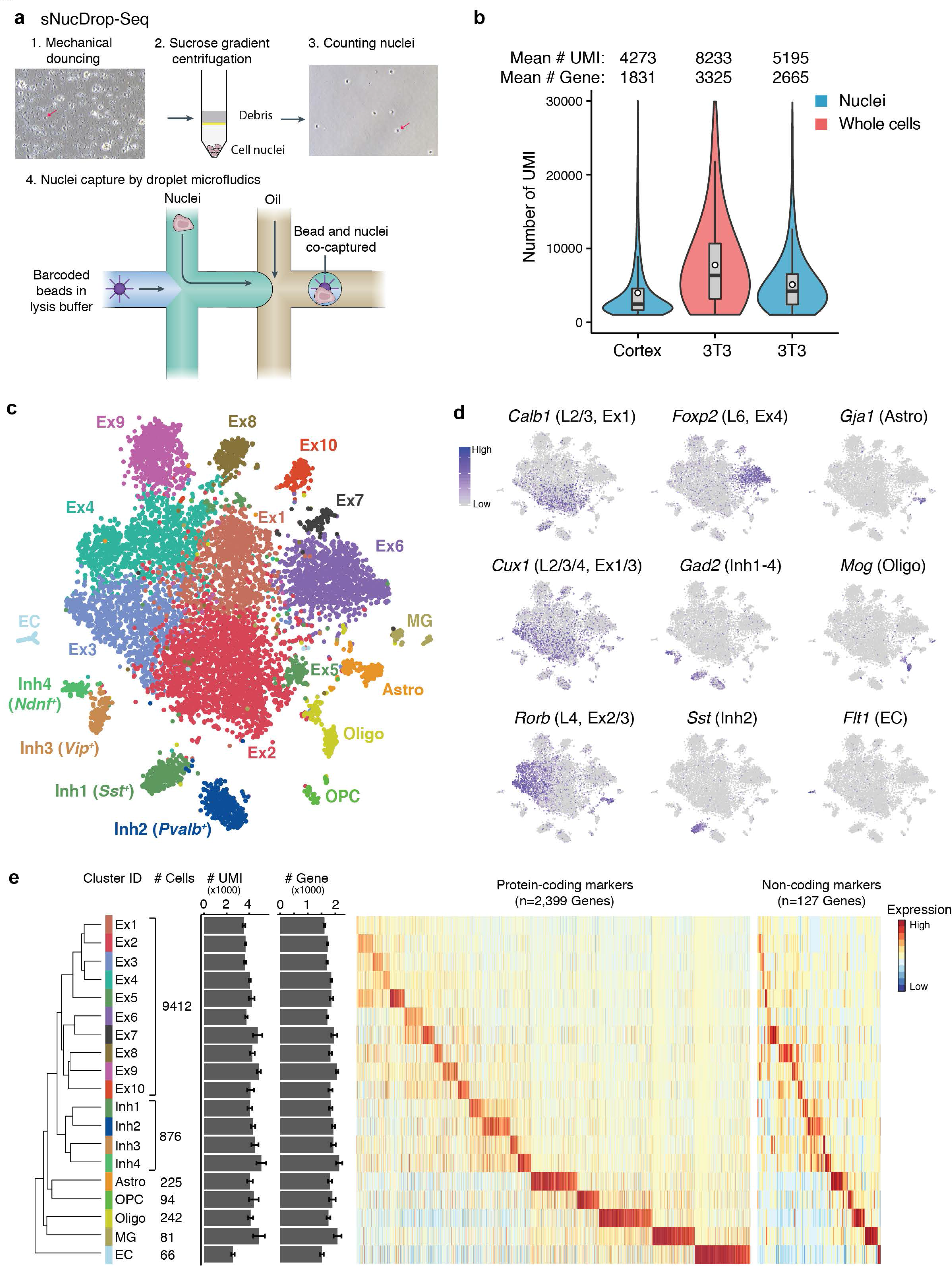
sNucDrop-Seq: a massively parallel single-nucleus RNA-Seq method. (a)Overview of sNucDrop-Seq. Step 1, dounce homogenization in lysis buffer is used to disrupt cellular membranes; Step2, nuclei are purified from cellular debris through sucrose gradient ultracentrifugation; Step3, quality and yield of nuclei is determined by hemocytometer count; Step4, nuclei and barcoded beads are co-encapsulated by an emulsion-droplet microfluidic device. Red arrows indicate representative nuclei before or after sucrose gradient centrifugation. (b)Violin plots illustrating number of transcripts (UMIs) detected by sNucDrop-Seq of nuclei isolated from mouse 3T3 cells (~23,000 reads per nucleus) and adult mouse cortex (~20,000 reads per nucleus) or by Drop-Seq of whole cells from 3T3 cells (~25,000 reads per cell). Center line: median; circle: mean; limits: first and third quartile; whiskers, ±1.5 IQR. Indicated on top are the number of cells or nuclei (>= 800 genes detected), mean number of UMIs per cells/nuclei, and mean number of genes per cells/nuclei. (c)Two-dimensional spectral t-stochastic neighborhood embedding (tSNE) plot of 11,283 nuclei isolated from adult mouse cortex, colored per density clustering and annotated according to known cell types. Ex, excitatory neurons; Inh, inhibitory neurons; Astro, astrocytes; OPC, oligodendrocyte precursor cells; Oligo, oligodendrocytes; MG, microglia; EC, endothelial cells. (d)Marker gene expression shown by re-coloring the tSNE plot. Shown is the same plot as Fig. 1c but with nuclei colored by the expression level of known cell type (e.g. Ex, Inh, Astro, Oligo, EC)- or cortical layer (L2/3/4/5/6)-specific marker genes. (e)Dendrogram illustrating relatedness of cell clusters, followed by (from left to right) cluster identification (ID), cell number per major cell type, UMIs per cluster (mean ± s.e.m.), number of genes detected per cluster (mean ± s.e.m.), heatmap showing protein-coding marker genes, and heatmap showing long non-coding RNA markers.

We next validated the specificity of sucrose gradient-assisted single-nucleus Drop-Seq (sNucDrop-Seq) with species-mixing experiments, using nuclei isolated from *in vitro* cultured mouse and human cells. This analysis indicates that the rate of co-encapsulation of multiple nuclei per droplet (~2.6%) is comparable to standard Drop-Seq (**Supplementary Fig. 3a**). To assess the sensitivity of sNucDrop-Seq, we performed shallow sequencing of cultured mouse 3T3 cells at either single-cell (with Drop-Seq: detecting on average 3,325 genes with ~25,000 reads per cell for 1,160 cells with >800 genes detected) or single-nucleus (with sNucDrop-Seq: detecting on average 2,665 genes with ~23,000 reads per nucleus for 1,984 nuclei with >800 genes detected) resolution (**Fig. 1b**). With standard Drop-Seq microfluidics devices and flow parameters, the throughput of sNucDrop-Seq (1.9%, 1,829 / 95,000 barcoded beads) is comparable to that of Drop-Seq (1.5%, 1,160 / 77,000 barcoded beads). Comparative analysis of Drop-Seq and sNucDrop-Seq reveals that mitochondria-derived RNAs (e.g. *mt-Nd1*, *mt-Nd2*) and nucleus-enriched long-noncoding RNAs (e.g. *Malat1*) were enriched in cytoplasmic and nuclear compartments, respectively (**Supplementary Fig. 3b**). Thus, integrating sucrose gradient centrifugation-based nuclei purification with the current Drop-Seq microfluidics device and workflow may support massively parallel single-nucleus RNA-Seq.

To demonstrate the utility of sNucDrop-Seq in studying complex adult tissues, we analyzed nuclei isolated from adult mouse cerebral cortex. The average expression profiles of single nuclei from two biologically independent replicates were well correlated (r=0.993; **Supplementary Fig. 3c**). Out of reads uniquely mapped to the genome (78.0% of all reads), 76.3% of reads were aligned to the expected strand of genic regions (25.3% exonic and 51.0% intronic), and the remaining 23.7% to intergenic regions or to the opposite strand of annotated genic regions. The relatively high proportion of intronic reads is similar to previous single-nucleus RNA-Seq study of human cortex (~48.7%) ^5^, reflecting the enrichment of nascent, pre-processed transcripts in the nucleus. Because most exonic (91.4%) and intronic (86.0%) reads were mapped to the expected strand of annotated transcripts, we retained both exonic and intronic reads for downstream analyses. After quality filtering, we retained 10,996 nuclei (~20,000 uniquely mapped reads per nucleus) from 13 animals, detecting, on average, 4,273 transcripts (unique molecular identifiers [UMIs]), and 1,831 genes per nucleus (**Fig. 1b**). After correcting for batch effects, we identified highly variable genes, and determined significant principal components (PC) with these variable genes. We then performed graph-based clustering and visualized distinct groups of cells using non-linear dimensionality reduction with spectral *t*-distributed stochastic neighbor embedding (tSNE) (**Methods**). This initial analysis segregated nuclei into 19 distinct clusters (**Fig. 1c**). Each cluster contains nuclei from multiple animals, indicating the transcriptional identities of these cell-type-specific clusters are reproducible across biological replicates (**Supplementary Fig. S4a**).

On the basis of known markers for major cell types, we identified 10 excitatory neuronal clusters (Ex 1-10; *Slc17a7^+^*), four inhibitory neuronal clusters (Inh 1-4; *Gad1^+^*), and five non-neuronal clusters (astrocytes [Astro; *Gja1^+^*], oligodendrocyte precursor cells [OPC; *Pdgfra^+^*], oligodendrocytes [oligo; *Mog^+^*], microglia [MG; *Ctss^+^*], and endothelial cells [EC; *Flt1^+^*]) (**Fig. 1c-d** and **Supplementary Fig. 4b**). We readily uncovered all major subtypes of GABAergic inhibitory neurons expressing known canonical markers: *Sst* (somatostatin; cluster Inh1), *Pvalb* (parvalbumin; cluster Inh2), *Vip* (vasoactive intestinal peptide; cluster Inh3) and *Ndnf* (neuron-derived neurotrophic factor; cluster Inh4) (**Supplementary Fig. 5a**). For glutamatergic excitatory neurons, hierarchical clustering grouped the ten clusters into two major groups (**Fig. 1e**), largely corresponding to their cortical layer positions, from superficial (cluster Ex1-5: L2/3 and L4) to deep (cluster Ex6-10: L5a/b and L6a/b) layers (**Fig. 1d** and **Supplementary Fig. 5**). Consistent with previous studies ^5, 13, 14^, we readily annotated anatomical location of each excitatory neuronal cluster *post-hoc* by its expression of known layer-specific marker genes (**Supplementary Fig. 6a-b**). In addition to protein-coding marker genes, we have also identified a list of long non-coding RNAs that are specifically expressed in distinct cell clusters (**Fig. 1e** and **Supplementary Fig. 5b**). For instance, *1700016P03Rik* is specifically detected in cluster Ex5, and this non-coding transcript acts mainly as a primary transcript encoding two neuronal activity-regulated microRNAs (*Mir212* and *Mir132*) ^15, 16^ (**Supplementary Fig. 7**), which is consistent with the enrichment of other activity-dependent genes (*Fos, Arc, Npas4*) in this excitatory neuronal cluster (**Supplementary Fig. 6a**), and raises the possibility that Ex5 is enriched of activated neurons (see below). The identification of both protein-coding and non-coding transcripts as cell-type-specific markers highlights the potential of sNucDrop-Seq in exploring the emerging role of non-coding RNAs at single-cell resolution *in vivo*.

Cortical interneurons are highly diverse in terms of morphology, connectivity and physiological properties ^17^. To further annotate these inhibitory neuronal subtypes, we performed sub-clustering on the 876 inhibitory neuronal nuclei in our dataset, identifying eight sub-clusters (cluster A-H in **Fig. 2a**). Unlike previous single-cell RNA-Seq analysis that employed pre-enrichment of cortical inhibitory neurons from transgenic mouse lines ^18^, sNucDrop-Seq samples the nuclei in proportion to cells’ abundance in their native environment, which provides a more accurate description of the cellular composition at the transcriptomic level. This analysis identified *Pvalb*-expressing subtypes (cluster D and E; *n*=359/876 nuclei, 41.0%) and *Sst*-expressing subtypes (cluster F, G, H; *n*=304/876 nuclei, 34.7%) as two major groups of cortical interneurons (**Fig. 2b-d**), in accordance with previous observations derived from *in situ* hybridization (ISH)- or immunostaining-based methods that *Pvalb-* and *Sst*-positive groups account for ~40% and ~30% of interneurons, respectively, in the neocortex ^19^. Beyond the major interneuron subtypes, we identified one *Ndnf*-expressing subtype (cluster A; *n*=84/876 nuclei), one *Vip*-expressing subtype (cluster B; *n*=74/876 nuclei), and one synuclein gamma (*Sncg*)-expressing subtype (cluster C; *n*=55/876 nuclei) (**Fig. 2b-d** and **Supplementary Fig. 8a**). On the basis of combinatorial expression of known marker genes associated with specific cortical layer and developmental origin, interneuron subtypes identified by sNucDrop-Seq parallel those identified from previous studies of mouse or human cortex ^5, 18^, revealing inhibitory neuronal heterogeneity in both cortical layer distribution (**Supplementary Fig. 8a-b**) and the developmental origin from subcortical regions of the medial or caudal ganglionic eminences (MGE or CGE) (**Fig. 2e**). Therefore, sNucDrop-Seq is able to resolve cellular heterogeneity and quantify cell-type composition at transcriptomic level with high sensitivity, including rare interneuron subtypes.

**Figure 2.**
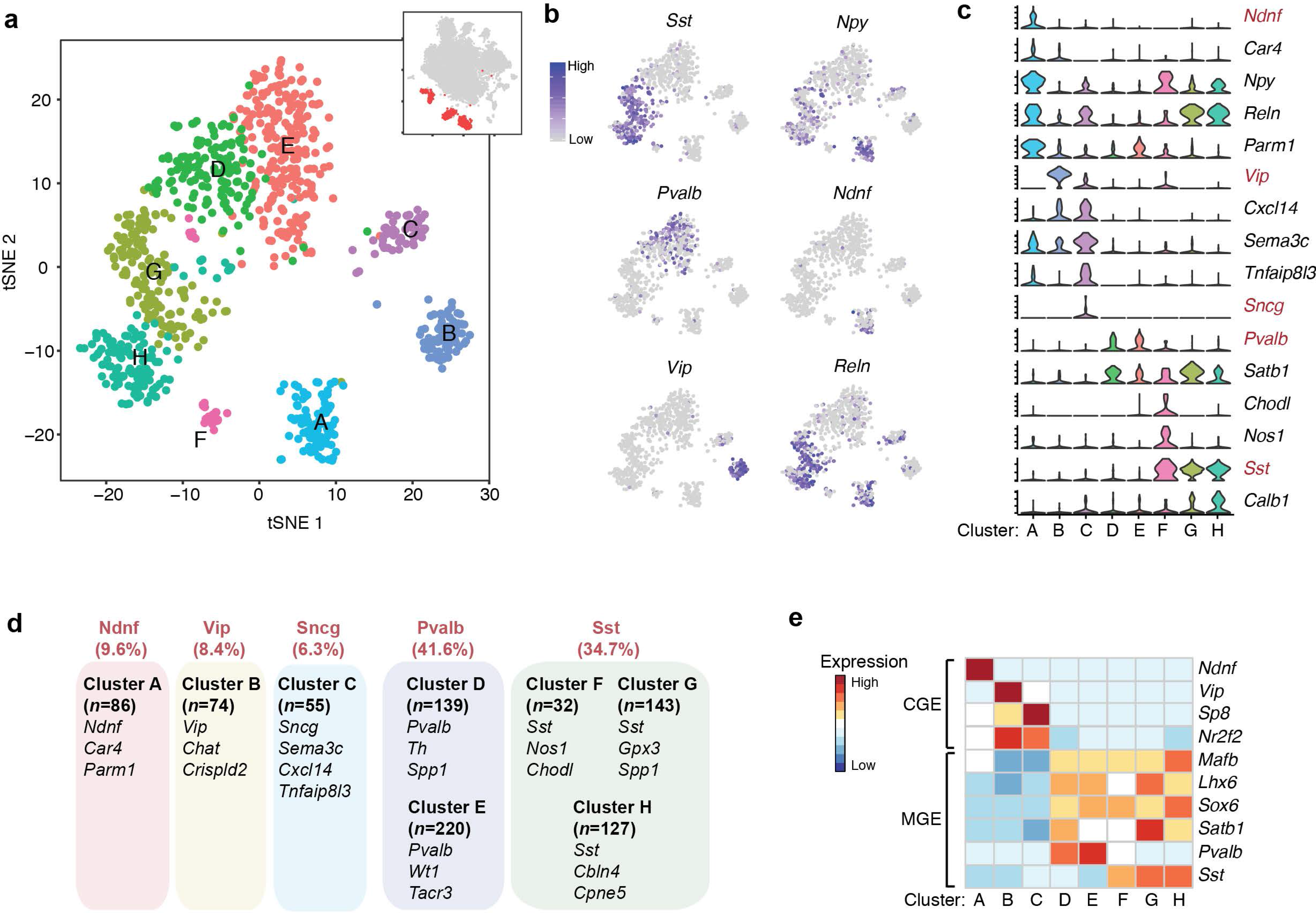
sNucDrop-Seq reveals inhibitory neuronal subtypes and composition. (a)Spectral tSNE plot of 876 inhibitory neurons, colored according to the results of sub-clustering (thumbnail: Fig. 1c). (b)Marker gene expression shown by re-coloring tSNE plot. Shown is the same plot as Fig. 2a but with nuclei colored by the expression level of known inhibitory neuronal subtype-specific marker genes. (c)Violin plots showing select marker gene expression for inhibitory neuronal subtypes. Five mutually exclusive subtype-specific marker genes are highlighted in red. (d)Summary of inhibitory neuronal subtypes identified by sNucDrop-Seq. GABAergic subtypes are grouped according to five major classes. Also shown are number of nuclei per subtype and representative marker genes for each subtype. (e)Heatmap showing select marker genes that distinguish inhibitory neurons originated from either CGE or MGE.

For glutamatergic neurons, unsupervised graph-based sub-clustering of two groups of excitatory neurons (upper layers versus lower layers) identified a total of 18 subtypes (Upper Ex 1-11 and Lower Ex 1-7; **Fig. 3a**). We associated each excitatory neuronal sub-cluster with a distinct combination of known markers indicative of their superficial-to-deep layer distribution (**Supplementary Fig. 9a**), capturing finer distinctions between closely related subtypes in each cortical layer, which is in high concordance with subtypes previously identified in human ^5^ and mouse ^14, 18^ cortices (**Fig. 3b** and **Supplementary Fig. 9b**). Beyond excitatory neuronal subtypes defined by cortical layer-specific markers, our analysis also resolved heterogeneity in neuronal activation states. In response to an activity-inducing experience, cortical excitatory neurons express a complex program of activity-dependent genes ^20^. Both upper-Ex3 (n=209; 3.1% of 6,770 nuclei in upper layer sub-clusters) and lower-Ex5 (n=213; 8.1% of 2,642 nuclei in lower layer sub-clusters) neurons are specifically associated with high-level expression of activity-dependent genes (**Fig. 3b** and **Supplementary Fig. 9c**), including immediately early genes (IEGs) such as *Fos*, *Arc*, and *Egr1* as well as other activity-regulated transcription factors (e.g. *Npas4*), genes encoding proteins that function at synapses (e.g. *Homer1*), and non-coding RNAs (e.g. *1700016P03Rik* that encodes Mir132). We determined the genes specifically enriched in upper-Ex3 (*n*=160 genes, as compared to other upper-Ex sub-clusters) or lower-Ex5 (*n*=134 genes, as compared to other lower-Ex sub-clusters) neurons (**Fig. 3c**). Transcriptional signatures identified in these two sub-populations are enriched for genes involved in the MAPK signaling pathway (e.g. *Dusp1*; adjusted *P*=2.67x10^-2^ for upper-Ex3 sub-cluster), as previously reported in low-throughput single-nucleus RNA-Seq analysis of *Fos*-positive nuclei isolated from the hippocampus of adult mice exposed to a novel environment ^2^. Together, these results demonstrate the utility of sNucDrop-Seq in the identification of transient transcriptional states, such as neuronal activation.

**Figure 3.**
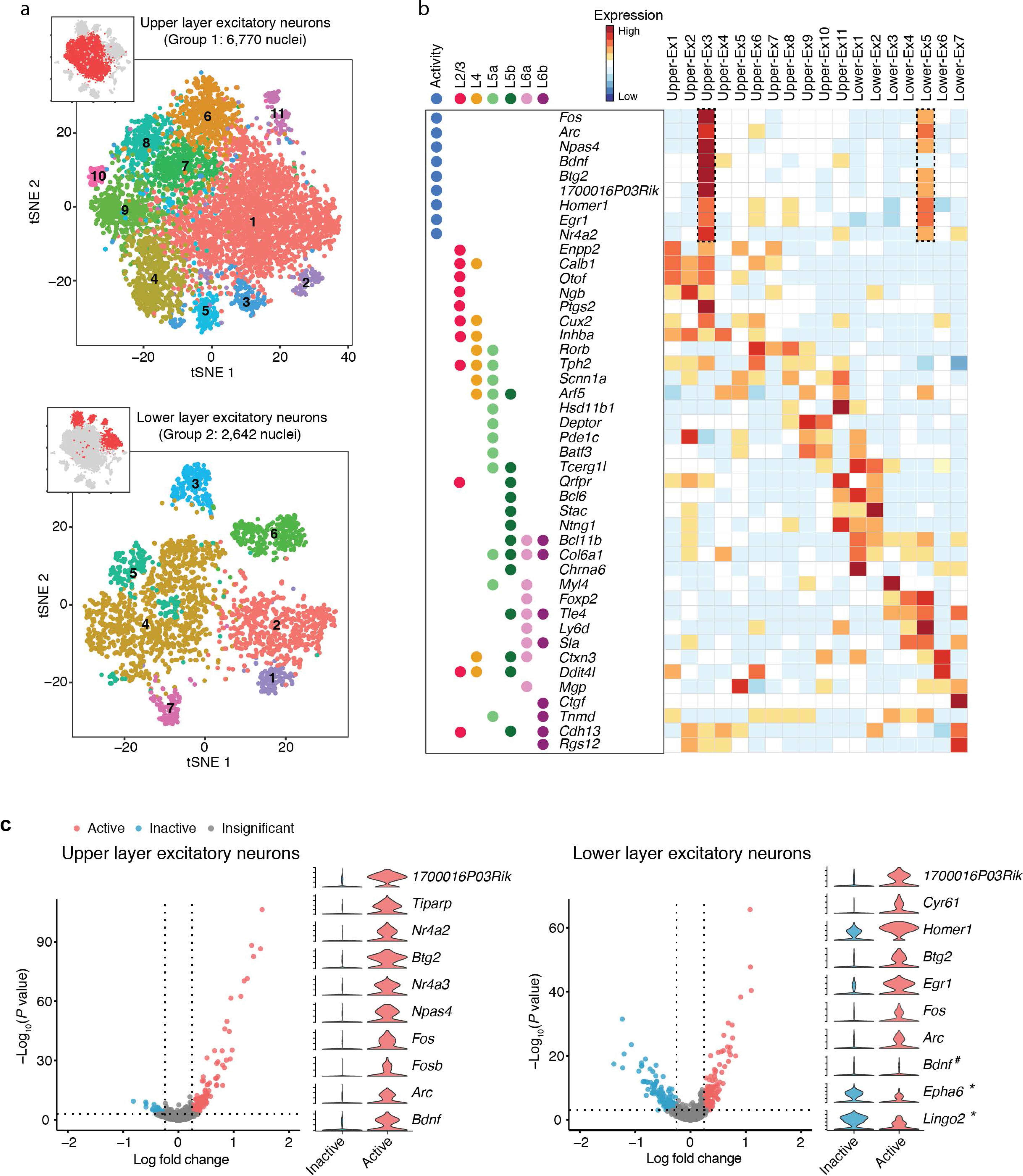
Excitatory neuronal subtypes resolve heterogeneity in cortical layer distribution and state of neuronal activity. (a)Spectral tSNE plots of 6,770 upper and 2,642 lower layer excitatory neurons, colored according the results of sub-clustering (thumbnails: Fig. 1c). (b)Heatmap for layer-specific markers and neuronal activity-regulated genes showing cortical layer identity (L2/3/, L4, L5a/b, L6a/b), excitatory subtypes, and activity-induced gene expression. (c)Differential expression between activated and inactivated excitatory neurons within upper (left) or lower (right) layer sub-clusters. Significant genes (red or blue), genes with p-values less than 0.001 and absolute natural log fold changes greater than 0.25. Violin plots showing select marker gene expression. #, denotes that the expression of *Bdnf* is not significantly different between active and inactive lower layer excitatory neurons. *, denotes that *Epha6* and *Lingo2* were expressed at significantly higher level in inactive lower layer excitatory neurons compared to active counterparts.

In conclusion, sNucDrop-Seq is a robust approach for massively parallel analysis of nuclear RNAs at single-cell resolution. Because intact nuclei isolation can potentially be accomplished by mechanical douncing and sucrose gradient ultracentrifugation in almost any primary tissue, including frozen archived human tissues, sNucDrop-Seq and similar approaches pave the way to systematically identify cell-types, reveal subtype composition, and dissect transient functional states such as activity-dependent transcription in complex mammalian tissues.

## ACKNOWLEDGEMENTS

Z.Z is supported by NIH grant R56MH111719. H.W. is supported by the National Human Genome Research Institute (R00HG007982).

## AUTHOR CONTRIBUTIONS

H.W. and Z.Z. conceived the project. H.W., P.H. and E.F. performed experiments and carried out data analysis. H.W. wrote the manuscript.

## Material and Methods

### Cell cultures and animals

Mouse NIH3T3 cells were purchased from ATCC (cat# CRL-1658) and were grown in Dulbecco’s Modified Eagle’s Medium (DMEM) (Life Technologies, cat# 11965084) supplemented with 10% fetal bovine serum (FBS) (Life Technologies, cat# 26140079) and 2 mM L-glutamine (Life Technologies, cat# 25030081) at 37°C in 5% CO_2_. The culture was passaged every 2-3 days using 0.05% Trypsin (Life Technologies, cat# 25300054). Human embryonic stem cells (H7) have been procured from WiCell (Madison, WI) and maintained at 37°C in 5% CO_2_ on growth factor-reduced Matrigel matrix (Corning, cat# 354230) coated six-well tissue culture plates. The six-well plates were coated with diluted (1:30) Matrigel matrix in DMEM/F12 (Life Technologies, cat# 11320033). H7 cells (between passage 40 and 70) were maintained in TeSR-E8 medium (Stem Cell Technologies, cat# 05940) and passaged every 5-6 days as small aggregates using an enzymatic digestion-free method (0.5 mM ETDA in D-PBS without CaCl_2_ and MgCl_2_).

Animal experiments were conducted in accordance with the ethical guidelines of the US National Institutes of Health and with the approval of the Institutional Animal Care and Use Committee of the University of Pennsylvania. All of the experiments described were performed using mice of C57BL/6 background.

### sNucDrop-Seq

#### Isolation and purification of nuclei

Mouse brains (postnatal 6 weeks) were rapidly resected on ice. Frozen cortices were flash frozen in liquid nitrogen for 2 minutes and subsequently kept at -80°C for 2 hours before nuclear isolation. 14 mL of sucrose cushion (1.8 M sucrose (CAS# 57-50-1, RNase & DNase free, ultra pure grade), 10 mM Tris-HCl pH 8.0 (Invitrogen, cat# 15568-025), 3 mM MgAc_2_ (CAS# 16674-78-5), protease inhibitor cocktail (Roche, cat# 11873580001)) was added to the bottom of centrifuge tubes (Beckman Coulter, cat# 326823). Using a glass homogenizer (Wheaton, cat# 357544), a freshly isolated or frozen mouse cortex sample was subjected to dounce homogenization (21 times with loose pestle followed by 7 times with tight pestle) in 12 mL of homogenization buffer (0.32M sucrose, 5 mM CaCl_2_ (CAS# 10043-52-4), 3mM MgAc_2_, 10 mM Tris-HCl pH 8.0, 0.1% Triton X-100 (CAS# 9002-93-1), 0.1 mM EDTA (Invitrogen, cat# 15575-020), protease inhibitor cocktail). For *in vitro* cultured cells, cell pellets (~5 million cells) were resuspended in homogenization buffer and dounced for 30 times with a loose pestle. Homogenates (~12 mL) were layered onto the sucrose cushion in the centrifuge tubes, and 10 mL of homogenization buffer was added atop of the homogenates. The tubes were then centrifuged in a Beckman Coulter L7-65 Ultracentrifuge at 25,000 rpm at 4°C for 2.5 hours using a Beckman Coulter SW28 swinging bucket rotor (Beckman Coulter, cat# 342207). The supernatant was carefully removed via aspiration. 1 mL of chilled DPBS with protease and RNase inhibitor (Lucigen, cat# 30281-2) was added to resuspend the nuclear pellet, and nuclei were subsequently transferred to a 1.5-mL tube. Nuclei were pelleted at 5,000 rpm for 10 min at 4°C, and then resuspended in 0.01% BSA (Sigma-Aldrich, cat# A8806-5G) in DPBS (Invitrogen, cat# 14190136). After resuspension, nuclei were filtered through a 40-μm cell strainer (Fisher Scientific, cat# 002087711), visually inspected for morphology and quality assurance, and counted using a Fuchs-Rosenthal counting chamber before droplet microfluidic encapsulation. For mouse cortices, we obtain 3.45 ± 2.00 x10^6^ nuclei, per round of isolation (based on 11 measurements). The nuclear isolation efficiency for *in vitro* cultured cells is ~84% (number of nuclei/number of input cells x 100).

#### Library preparation and sequencing

The nuclear suspension was diluted to a concentration of 100 nuclei/μL in DPBS containing 0.01% BSA. Approximately 1.25 mL of this single-nucleus suspension was loaded for each sNucDrop-Seq run. The single-nucleus suspension was then co-encapsulated with barcoded beads (ChemGenes, cat# MACOSKO-2011) using an Aquapel-coated PDMS microfluidic device (uFluidix), connected to syringe pumps (KD Scientific) via polyethylene tubing with an inner diameter of 0.38 mm (Scientific Commodities, cat# BB31695-PE/2). Barcoded beads were resuspended in lysis buffer (200 mM Tris-HCl pH8.0, 20 mM EDTA, 6% Ficoll PM-400 (GE Healthcare/Fisher Scientific, 45-001-745), 0.2% Sarkosyl (Sigma-Aldrich, cat# L7414-50mL), and 50 mM DTT (Fermentas, cat# R0862; freshly made on the day of run)) at a concentration of 120 beads/μL. The flow rates for cells and beads were set to 4,000 μL/hour, while QX200 droplet generation oil (Bio-rad, cat# 186-4006) was run at 15,000 μL/hour. A typical run lasts ~20 min. Droplet breakage with Perfluoro-1-octanol (Sigma-Aldrich, cat# 370533-25G), reverse transcription and exonuclease I treatment were performed, as previously described, with minor modifications ^7^. Specifically, up to 120,000 beads, 200 μL of reverse transcription (RT) mix (1x Maxima RT buffer, 4% Ficoll PM-400, 1 mM dNTPs (Clontech, cat# 639125), 1 U/μL RNase inhibitor (Lucigen, cat# 30281-2), 2.5 μM Template Switch Oligo (TSO; AAGCAGTGGTATCAACGCAGAGTGAATrGrGrG) _7_, and 10 U/μL Maxima H Minus Reverse Transcriptase (Life Technologies, cat# EP0753)) were added. The RT reaction was incubated at room temperature for 30 minutes, followed by incubation at 42°C for 150 minutes. To determine an optimal number of PCR cycles for amplification of cDNA, an aliquot of 6,000 beads (corresponding to ~100 nuclei) was amplified by PCR in a volume of 50 μL (25 μL of 2x KAPA HiFi hotstart readymix (KAPA biosystems, cat# KK2602), 0.4 μL of 100 μM TSO-PCR primer (AAGCAGTGGTATCAACGCAGAGT) ^7^, 24.6 μL of nuclease-free water) with the following thermal cycling parameter (95°C for 3 min; 4 cycles of 98°C for 20 sec, 65°C for 45 sec, 72°C for 3 min; 9 cycles of 98°C for 20 sec, 67°C for 45 sec, 72°C for 3 min; 72°C for 5 min, hold at 4°C). After two rounds of purification with 0.6x SPRISelect beads (Beckman Coulter, cat# B23318), amplified cDNA was eluted with 10 μL of water. 10% of amplified cDNA was used to perform real-time PCR analysis (1 μL of purified cDNA, 0.2 μL of 25 μM TSO-PCR primer, 5 μL of 2x KAPA FAST qPCR readymix, and 3.8 μL of water) to determine the additional number of PCR cycles needed for optimal cDNA amplification (Applied Biosystems QuantStudio 7 Flex). We then prepared PCR reactions per total number of barcoded beads collected for each sNucDrop-Seq run, adding 6,000 beads per PCR tube, and ran the aforementioned program to enrich the cDNA for 4 + 10 to 12 cycles. We then tagmented cDNA using the Nextera XT DNA sample preparation kit (Illumina, cat# FC-131-1096), starting with 600 pg of cDNA pooled in equal amounts, from all PCR reactions for a given run. Following cDNA tagmentation, we further amplified the library with 12 enrichment cycles using the Illumina Nextera XT i7 primers along with the P5-TSO hybrid primer (AATGATACGGCGACCACCGAGATCTACACGCCTGTCCGCGGAAGCAGTGGTATCAACGCAGAGT*A*C) ^7^. After quality control analysis using a Bioanalyzer (Agilent), libraries were sequenced on an Illumina NextSeq 500 instrument using the 75-cycle High Output v2 Kit (Illumina cat# FC-404-2005). We loaded the library at 1.9 pM and provided Custom Read1 Primer (GCCTGTCCGCGGAAGCAGTGGTATCAACGCAG AGTAC) at 0.3 μM in position 7 of the reagent cartridge. The sequencing configuration was 20 bp (Read1), 8 bp (Index1), and 50 or 60 bp (Read2). In total, 13 mouse cortex samples were analyzed with sNucDrop-Seq in four sequencing runs.

### Single-cell RNA-Seq library preparation using Drop-Seq and 10x Genomics platforms

Drop-Seq was performed as previously described, with default settings^7^. The cell suspension was diluted to 100 cells/μL with DPBS containing 0.01% BSA and 1 mL cell suspension was loaded for each Drop-seq run. After cell capture, reverse transcription, exonuclease treatment, cDNA amplification and tagmentation, libraries were diluted to 1.9 pmol, and equal amounts of distinctively indexed libraries were mixed and subjected to 75 cycles of paired-end sequencing on Illumina NextSeq 500 sequencer. 10x Genomics single-cell 3’ libraries were constructed, as previously described ^9^, with recommended settings using Chromium single cell 3’ v2 reagent kits.

### Data analysis of sNucDrop-Seq

#### Preprocessing of sNucDrop-Seq data

Paired-end sequencing reads of Single-nucleus RNA-seq were processed using publicly available the Drop-Seq Tools v1.12 software _7_ with some modifications. Briefly, each mRNA read (read2) was tagged with the cell barcode (bases 1 to 12 of read 1) and unique molecular identifier (UMI, bases 13 to 20 of read 1), trimmed of sequencing adaptors and poly-A sequences, and aligned using STAR v 2.5.2a to the mouse (mm10, Gencode release vM13) or a concatenation of the mouse and human (for the species-mixing experiment) reference genome assembly. Because a substantial proportion (~50%) of reads derived from nuclear transcriptomes of mouse cortices were mapped to the intronic regions, the intronic reads were retained for downstream analysis. A custom Perl script was implemented in the Drop-Seq Tools pipeline to retrieve both exonic and intronic reads mapped to predicted strands of annotated genes. Uniquely mapped reads were grouped by cell barcodes. Cell barcodes were corrected for possible bead synthesis errors, using the *DetectBeadSynthesisErrors* program from the Drop-Seq Tools v1.12 software. To generate digital expression matrix, a list of UMIs in each gene (as rows), within each cell (as columns), was assembled, and UMIs within ED = 1 were merged together. The total number of unique UMI sequences was counted, and this number was reported as the number of transcripts of that gene for a given cell.

#### Cell clustering and marker gene identification

Raw digital expression matrices were combined and loaded into the R package Seurat. For normalization, UMI counts for all nuclei were scaled by library size (total UMI counts), multiplied by 10,000 and transformed to log space. Only genes found to be expressing in >10 cells were retained. Nuclei with a relatively high percentage of UMIs mapped to mitochondrial genes (>=0.1) were discarded. Moreover, nuclei with fewer than 800 or more than 6,000 detected genes were omitted, resulting in 11,471 nuclei that pass filter.

Before clustering, batch effects from multiple animals were regressed out using the function *RegressOut* in R package Seurat. The highly variable genes were identified using the function *MeanVarPlot* with the parameters: x.low.cutoff = 0.0125, x.high.cutoff = 3 and y.cutoff = 0.8, resulting in an output of 2,178 highly variable genes. The expression level of highly variable genes in the nuclei was scaled and centered along each gene, and was conducted to principal component analysis. We then used two methods to assess the number of PCs to be utilized in downstream analysis: 1) The cumulative standard deviations accounted for by each PC were plotted using the function *PCElbowPlot* in Seurat to identify the ‘knee’ point at a PC number after which successive PCs explain diminishing degrees of variance, and 2) the significance for each gene’s association with each PC was accessed by the function *JackStraw* in Seurat. Based on these two methods, we selected first 20 PCs for two-dimensional t-distributed stochastic neighbor embedding (tSNE), implemented by the Seurat software with the default parameters. Based on the tSNE map, twenty-one clusters were identified using the function *FindCluster* in Seurat with the resolution parameter set to 1.0. Clusters that co-express both non-neuronal and neuron markers, representing cell doublets, were removed. As a result, we were able to assign 11,283 nuclei (98.4% of our data) into 19 cell type clusters.

To identify the marker genes, differential expression analysis was performed by the function *FindAllMarkers* in Seurat with likelihood-ratio test. Differentially expressed genes that were expressed at least in 30% cells within the cluster and with a fold change more than 0.25 (log scale) were considered to be marker genes. In total, 2,399 protein-coding genes and 127 long non-coding RNAs were identified for 19 clusters. For the marker genes, average gene expression for each cluster was determined, and Euclidean distances between all pairs was calculated. This dataset was used as input for complete linkage hierarchical clustering and dendrogram assembly. To generate a heatmap of marker genes across clusters, the average expression level of marker genes within each cluster were calculated. For each cluster, the average expression was centered and scaled by each gene. Next, the *heatmap.2* function in the R gplots package was used to generate the heatmap, and the parameter “dendrogram” was set to "column" to show the cluster dendrogram.

#### Sub-clustering

Excitatory neuronal nuclei from upper layers (cluster Ex1, Ex2, Ex3, Ex4, and Ex5) and deep layers (cluster Ex6, Ex7, Ex8, Ex9, and Ex10) were first combined, then sub-clustered using the same strategy described above, respectively. To correct for potential over-clustering, we merged the clusters showing high similarity. Briefly, pairwise comparison was conducted to identify the differentially expressed genes using the function *FindAllMarkers* in Seurat, with likelihood-ratio test. We merged the clusters showing <5 genes with an average expression difference greater than 2-fold between clusters. Specifically, we identified 2,218 highly variable genes associated with 12 PCs in 6,770 upper layer nuclei, which were further assigned into 11 sub-clusters, while 2,642 lower layer nuclei, containing 1,969 highly variable genes along with 9 PCs, were sub-clustered into 7 groups. After filtering out 138 unassigned nuclei, we grouped 9,412 nuclei (98.6 % of excitatory nuclei) into 17 layer-specific sub-clusters. For a comparison between active and inactive neurons, differentially expressed genes were identified using a likelihood ratio test with the p-value threshold set to 0.001.

For inhibitory neurons, nuclei from clusters Inh1, Inh2, Inh3 and Inh4 were combined and loaded into Seurat for sub-clustering. We identified 2,337 highly variable genes in 1,025 nuclei that were subjected to PCA analysis. First 10 PCs were selected for tSNE analysis and sub-clustering. After filtering out 149 unassigned nuclei, we identified 8 clusters, made up of 876 nuclei.

### GO enrichment analysis

To identify functional categories associated with genes enriched in activated neurons, the GO annotations were downloaded from the Ensembl Biomart database and KEGG annotations were retrieved by KEGG API. An enrichment analysis was performed via a hypergeometric test. The p-value was calculated using the following formula:

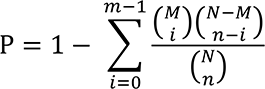

where N is the total number of genes, n is the total number of marker genes, M is the number of genes annotated to a certain GO term or KEGG pathway, and i is the number of marker genes annotated to a certain GO term or KEGG pathway. The p-value was corrected by function p.adjust with a false discovery rate (FDR) correction in R. GO terms or KEGG pathways with a FDR below 0.05 were considered enriched. All statistical calculations were performed in R.

## Supplementary Figures

**Supplementary Figure 1.**
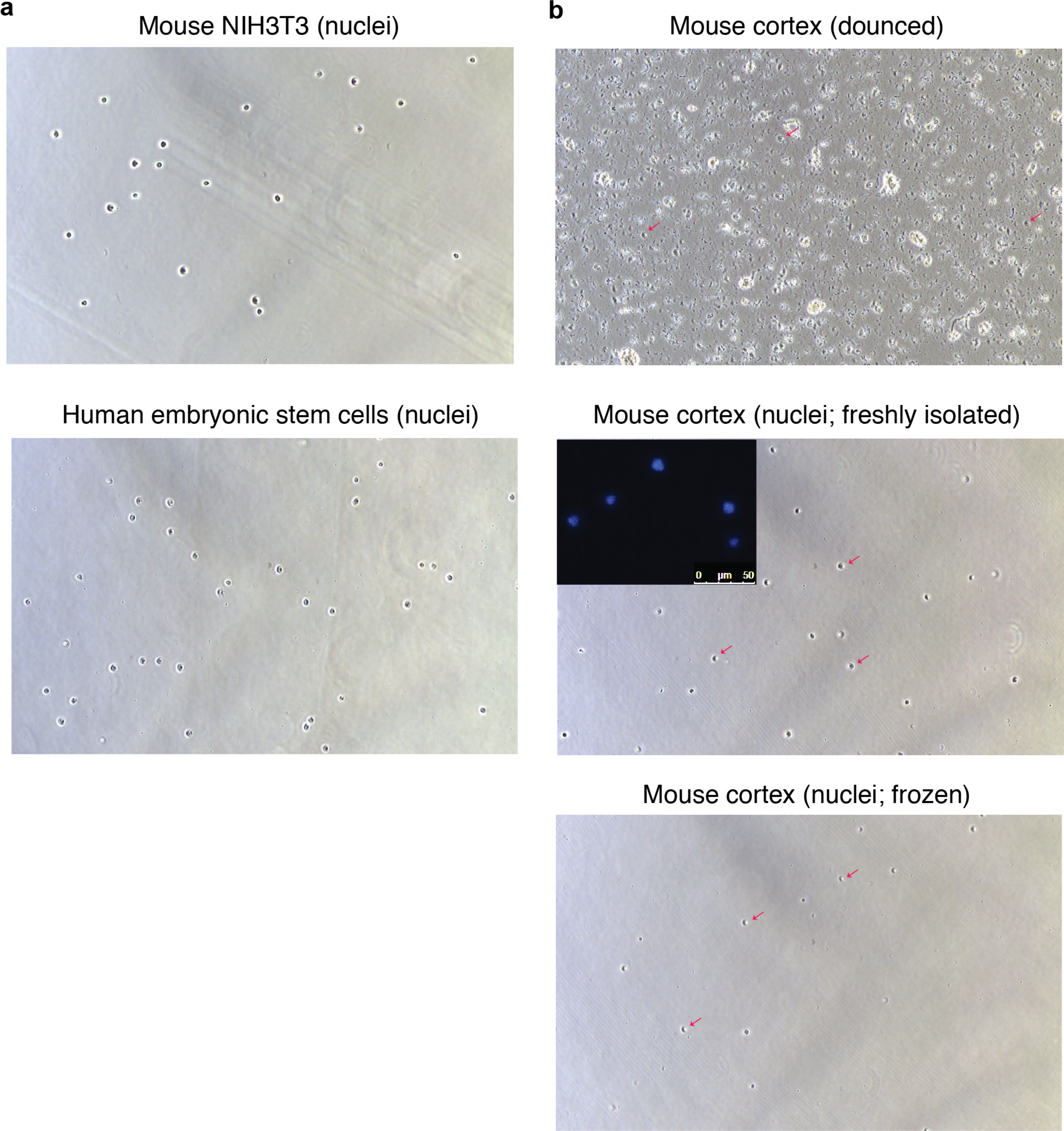
Quality control of nuclei isolation. **(a)** After sucrose gradient centrifugation and filtering through cell mesh, mouse (top, NIH3T3) and human (bottom, embryonic stem cells (ESCs)) nuclei were visualized by phase-contrast microscopy (10x). **(b)** After dounce homogenization, mouse cortical nuclei were visualized by phase-contrast microscopy (10x) before (top) or after (bottom) sucrose gradient centrifugation. Red arrows indicate nuclei before or after sucrose gradient centrifugation. The inlet indicates fluorescent image of purified nuclei stained with DNA intercalating dye Hoechst 33342 (10 ng/µL). Scale bar, 50 µm.

**Supplementary Figure 2.**
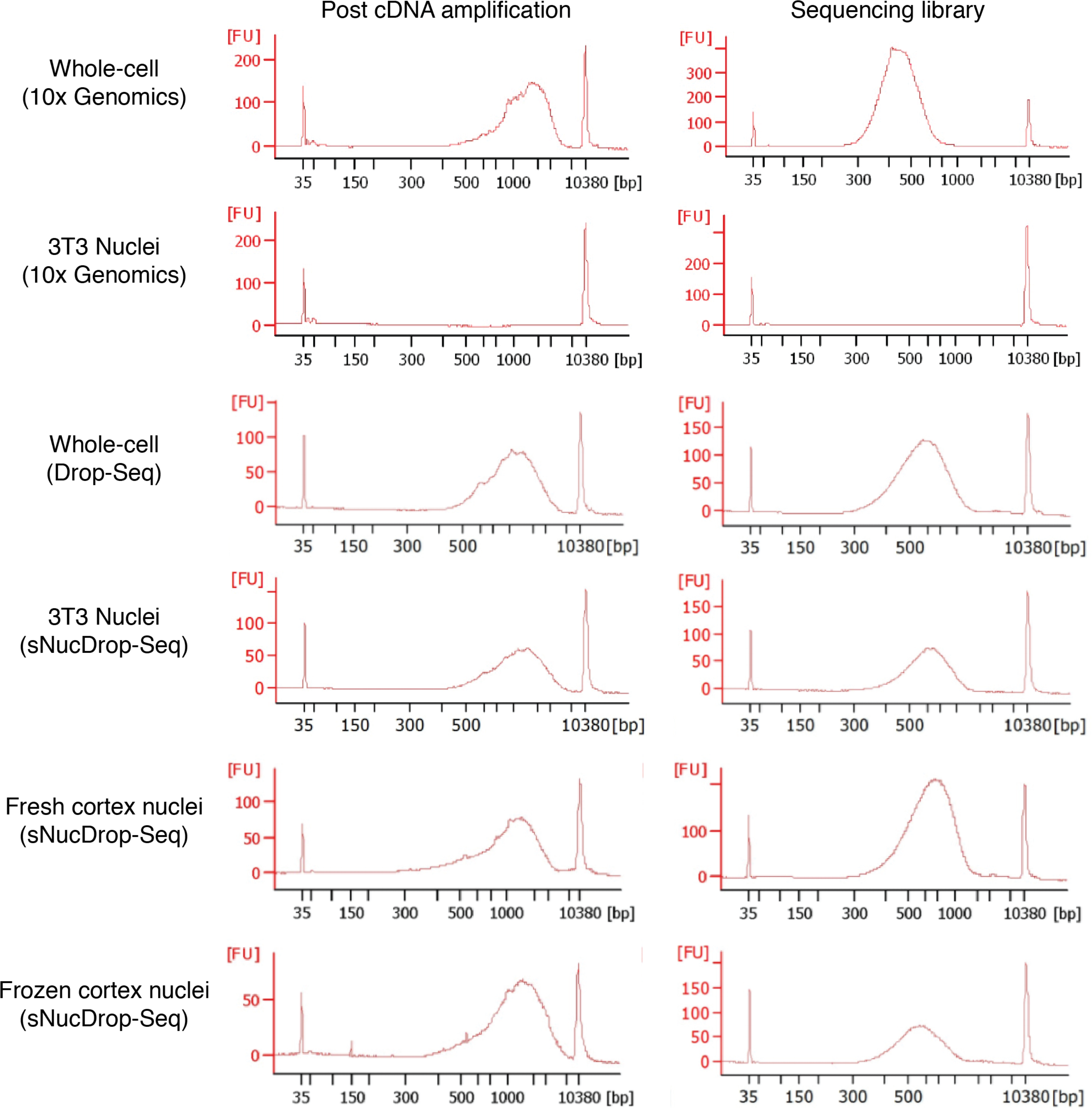
Testing different microfluidics platforms and library preparation workflows for single-nucleus RNA-Seq. Bioanalzyer electropherogram of amplified cDNA (left) and final sequencing library (right) shown for samples prepared from whole-cell or nuclei by different platforms (10x Genomics platform or Drop-Seq/sNucDrop-Seq). FU, fluorescence units.

**Supplementary Figure 3.**
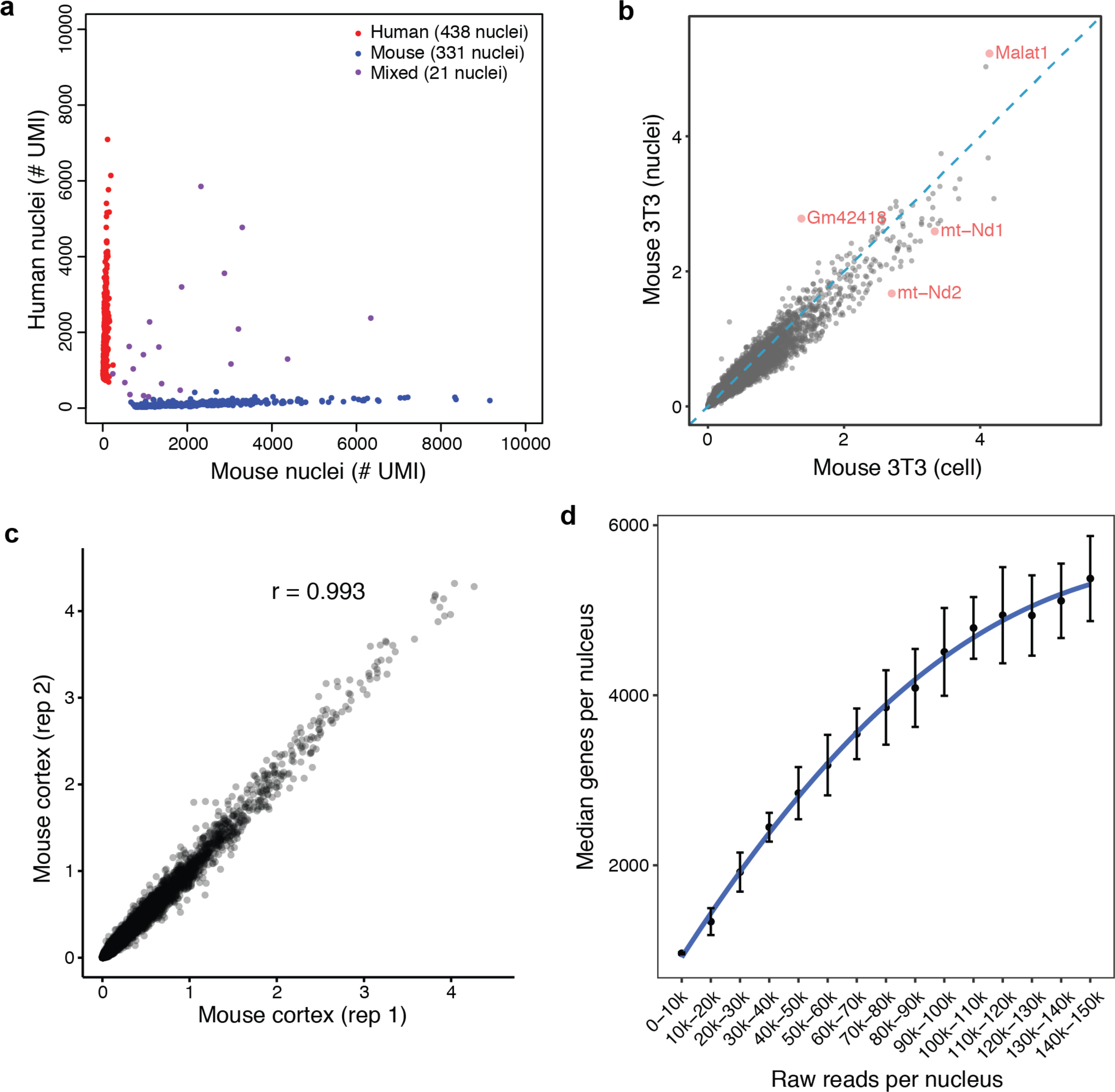
Specificity and performance of sNucDrop-Seq. **(a)** Multi-species nuclei-mixing experiment measures sNucDrop-Seq specificity, by sequencing a mix of human (ESCs) and mouse (NIH3T3) nuclei. Scatter plot shows the number of transcripts (UMIs) associated with annotated human (y-axis) or mouse (x-axis) transcripts for each nucleus (dot). Nuclei with >80% human transcripts are labeled as human (red), and nuclei with >80% mouse transcripts are labeled as mouse (blue). Nuclei with a relatively high percentage of both human and mouse transcripts are labeled as mixed (purple). Of the 790 nuclei that passed quality filter (>800 UMIs), 21 (2.66%) had a mixed phenotype. **(b)** Scatter plot comparing the average expression levels detected in single NIH3T3 nuclei (y-axis, by sNucDrop-Seq) and cells (x-axis, by Drop-Seq). Red dots mark representative genes preferentially enriched in either nuclei or cytoplasm (cell). For comparison, digital expression matrices of cell and nuclei were first combined, and UMI counts were then scaled by library size (total UMI counts per cell or nuclei), multiplied by 10,000 and natural log transformed. Only cells or nuclei that expressed >800 genes were retained for analysis. For each gene, the average normalized expression level were calculated as log (normalized UMI counts + 1). **(c)** Scatter plot showing the high correlation of average expression levels [log (normalized UMI counts + 1)] between two biological replicates of sNucDrop-Seq analysis of mouse cortex. **(d)** Median number of genes detected per nucleus at different raw reads per nucleus. Data from two independent experiments were included, mean±s.e.m.

**Supplementary Figure 4.**
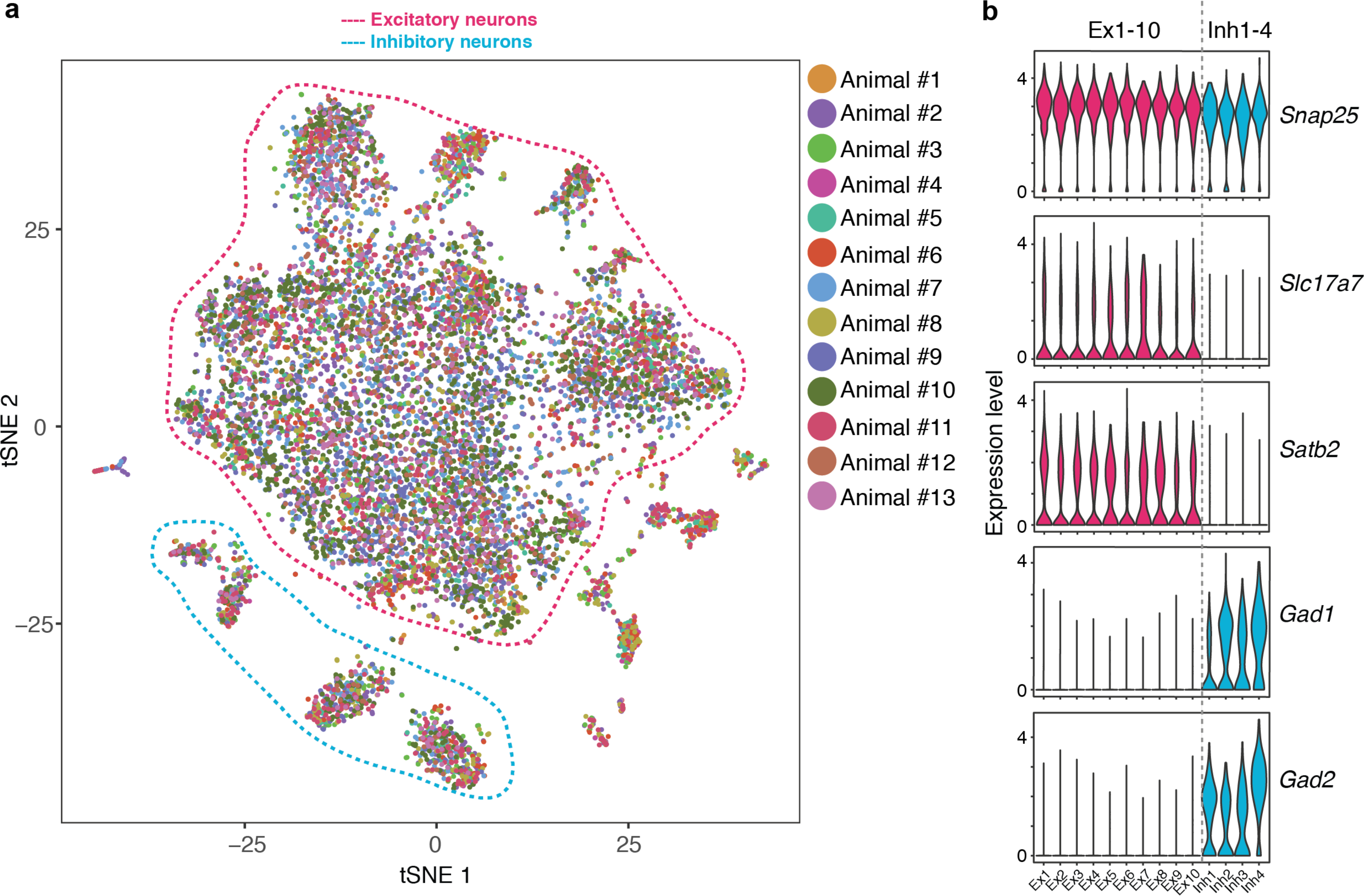
Cluster composition and neuronal marker gene expression. **(a)** The tSNE plot (the same plot as in Fig. 1c) with all nuclei colored according to animal identity. Clusters corresponding to excitatory (red dashed line) and inhibitory (blue dashed line) neurons are grouped together. **(b)** Violin plot illustrating the expression of pan-neuronal (*Snap2*5), excitatory neuronal (*Slc17a7, Satb2*) and inhibitory neuronal (*Gad1, Gad2*) markers for excitatory (red, Ex1-10) and inhibitory (blue, Inh 1-4) neuronal clusters.

**Supplementary Figure 5.**
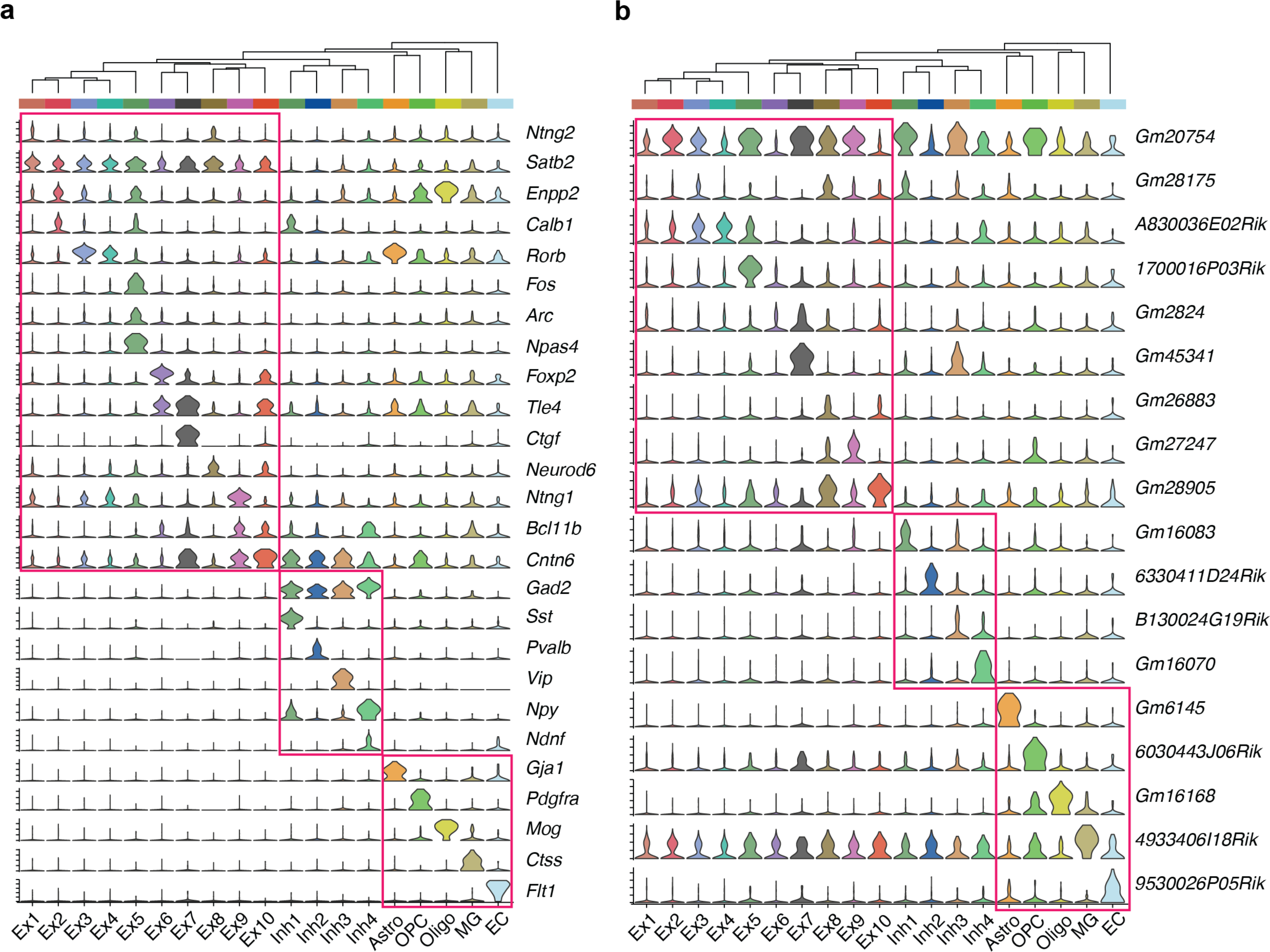
Protein-coding and noncoding marker gene expression for neuronal and non-neuronal cell clusters. Violin plots illustrating select protein-coding (**a**) and non-coding (**b**) marker gene expression for excitatory neuronal (Ex1-10), inhibitory neuronal (Inh 1-4) and non-neuronal (Astro, OPC, Oligo, MG, EC) cell clusters.

**Supplementary Figure 6.**
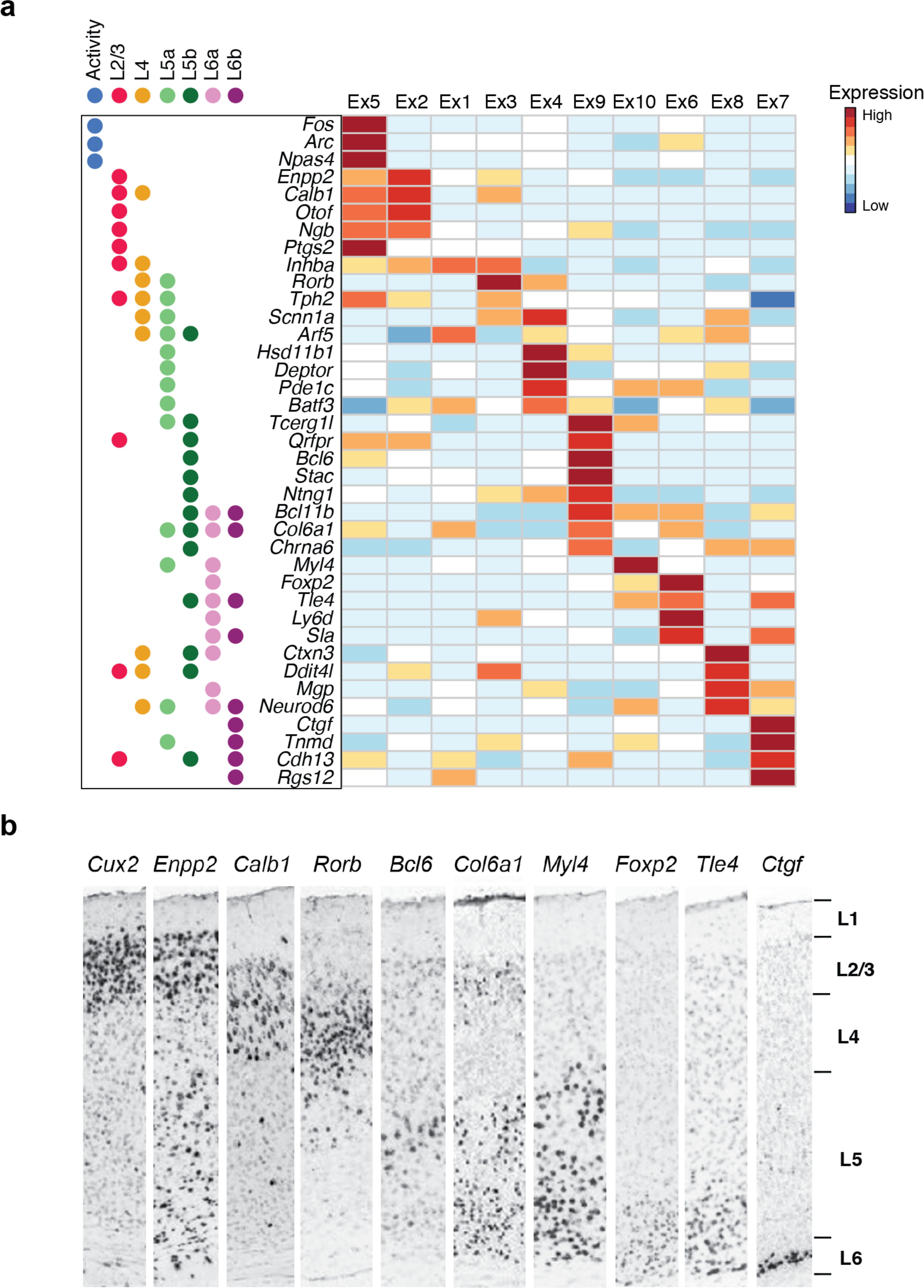
Cortical layer identity and activity-dependent transcriptional states of excitatory neurons. **(a)** Heatmap showing layer-specific markers (L2/3/, L4, L5a/b, L6a/b) and neuronal activity-regulated gene expression in excitatory neuronal clusters (Ex 1-10 identified Fig. 1c). **(b)** RNA in situ hybridization (ISH) showing layer-specific expression of selected markers in the mouse adult cortex (postnatal day 56, Allen Brain Atlas).

**Supplementary Figure 7.**
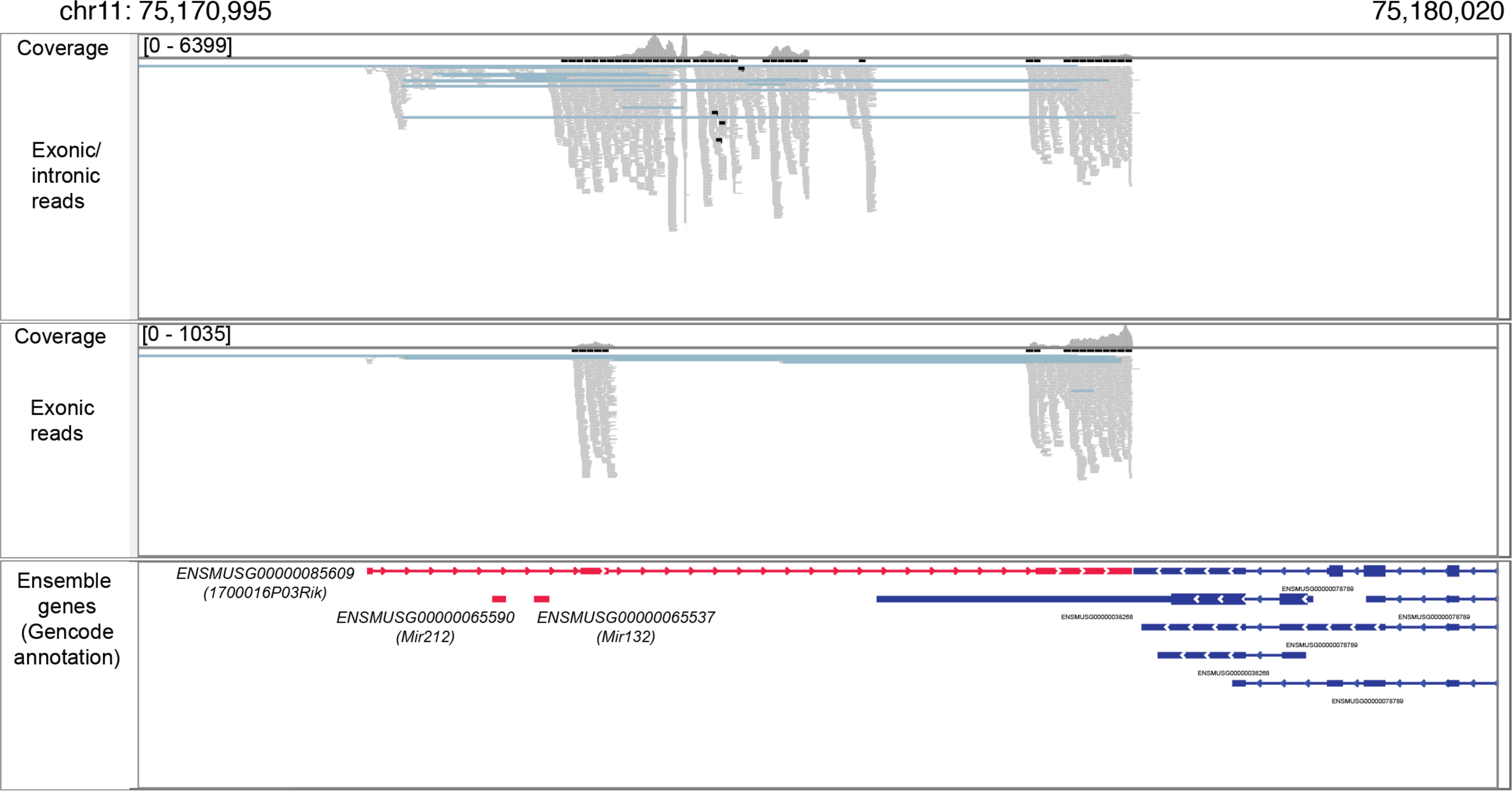
Distribution of exonic and intronic reads mapped to an activity-regulated non-coding transcript. Genome browser view (build: mm10) of exonic and intronic reads (from sNucDrop-Seq of mouse cortex) mapped to *1700016P03Rik* (highlighted in red), a neuronal activity-induced non-coding transcript that encodes two microRNAs (Mir212 and Mir132).

**Supplementary Figure 8.**
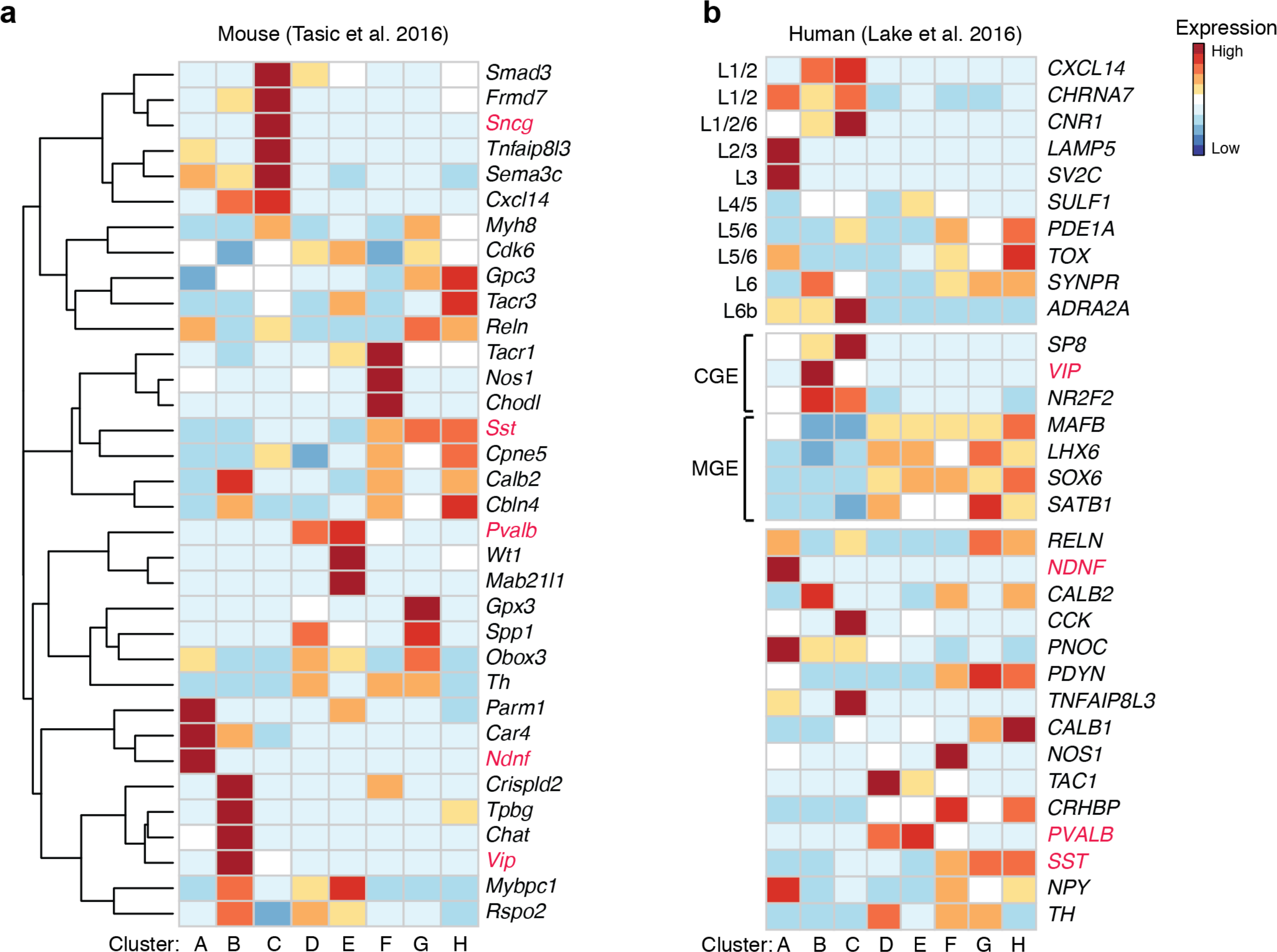
Marker gene expression for inhibitory neuronal subtypes. Heatmap illustrating select mouse (**a**) and human (**b**) marker gene expression for cortical inhibitory neuronal sub-populations (cluster A-H) identified in Fig. 2a. The mouse marker gene list is derived from Tasic et al. (2016) ^1^. The human marker gene list is derived from Lake et al. (2016) ^2^. Five mutually exclusive subtype-specific marker genes are highlighted in red. CGE, caudal ganglionic eminences; MGE, medial ganglionic eminences.

**Supplementary Figure 9.**
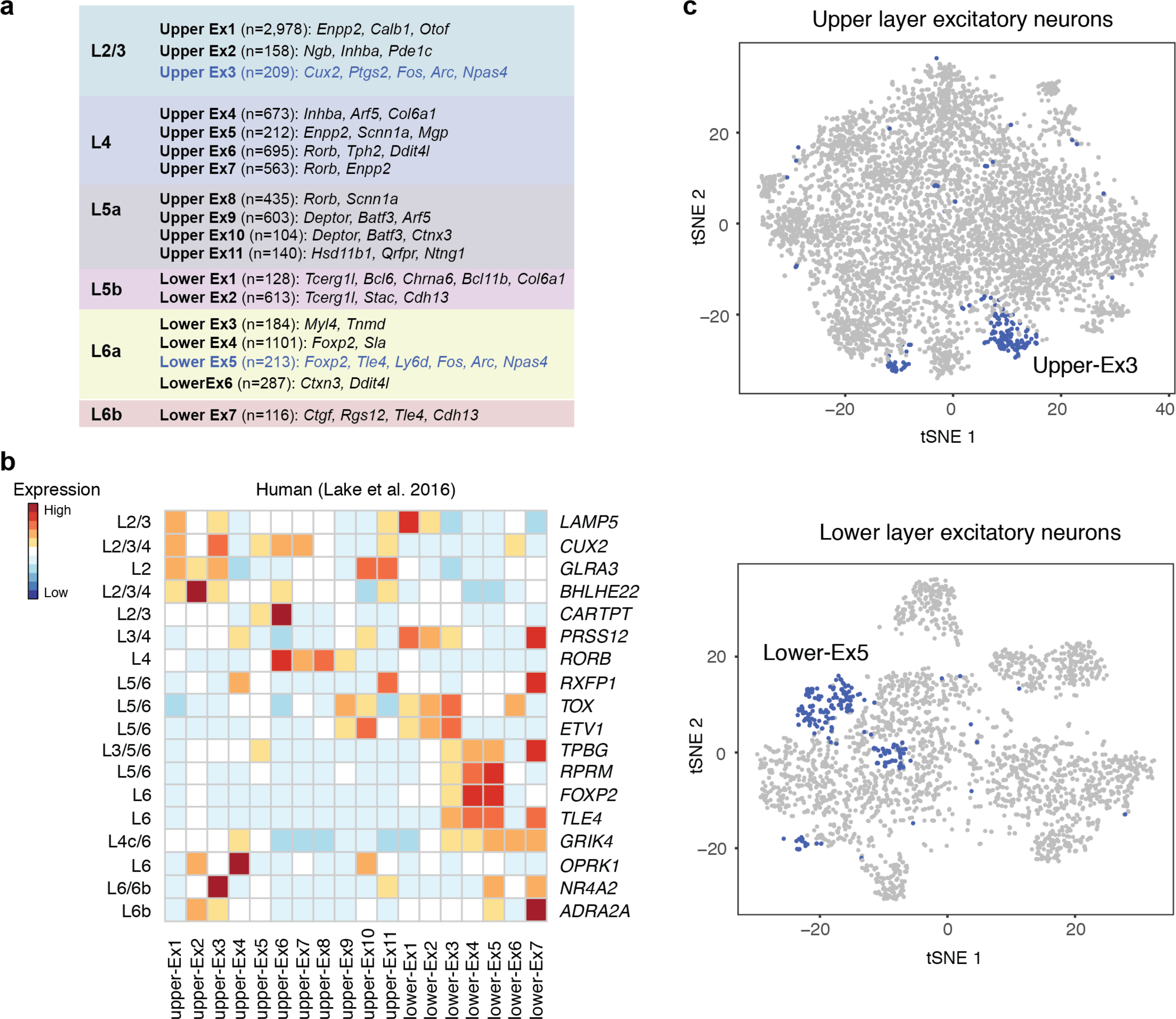
Cortical layer- and neuronal activity-dependent marker gene expression for excitatory neuronal subtypes. **(a)** Summary of excitatory neuronal subtypes identified by sNucDrop-Seq. Glutamatergic neuronal subtypes are grouped according to cortical layer distribution. Also shown are number of nuclei per subtype and representative marker genes for each subtype. **(b)** Heatmap showing select human marker gene expression for cortical excitatory neuronal sub-populations identified in Fig. 3a. The human marker gene list is derived from Lake et al. (2016) ^2^. **(c)** Spectral tSNE plots (the same plots in Fig. 3a) highlighting activated excitatory neurons in upper (top, upper-Ex3) and lower (bottom, lower-Ex5) layer sub-clusters.

## References

1. Tanay, A. & Regev, A. Scaling single-cell genomics from phenomenology to mechanism. Nature 541, 331–338 (2017).

2. Lacar, B. et al. Nuclear RNA-seq of single neurons reveals molecular signatures of activation. Nat Commun 7, 11022 (2016).

3. White, R.B., Bierinx, A.S., Gnocchi, V.F. & Zammit, P.S. Dynamics of muscle fibre growth during postnatal mouse development. BMC Dev Biol 10, 21 (2010).

4. Habib, N. et al. Div-Seq: Single-nucleus RNA-Seq reveals dynamics of rare adult newborn neurons. Science 353, 925–928 (2016).

5. Lake, B.B. et al. Neuronal subtypes and diversity revealed by single-nucleus RNA sequencing of the human brain. Science 352, 1586–1590 (2016).

6. Zeng, W. et al. Single-nucleus RNA-seq of differentiating human myoblasts reveals the extent of fate heterogeneity. Nucleic Acids Res 44, e158 (2016).

7. Macosko, E.Z. et al. Highly Parallel Genome-wide Expression Profiling of Individual Cells Using Nanoliter Droplets. Cell 161, 1202–1214 (2015).

8. Klein, A.M. et al. Droplet barcoding for single-cell transcriptomics applied to embryonic stem cells. Cell 161, 1187–1201 (2015).

9. Zheng, G.X. et al. Massively parallel digital transcriptional profiling of single cells. Nat Commun 8, 14049 (2017).

10. Jiang, Y., Matevossian, A., Huang, H.S., Straubhaar, J. & Akbarian, S. Isolation of neuronal chromatin from brain tissue. BMC Neurosci 9, 42 (2008).

11. Mo, A. et al. Epigenomic Signatures of Neuronal Diversity in the Mammalian Brain. Neuron 86, 1369–1384 (2015).

12. Lister, R. et al. Global epigenomic reconfiguration during Mammalian brain development. Science 341, 1237905 (2013).

13. Madisen, L. et al. Transgenic mice for intersectional targeting of neural sensors and effectors with high specificity and performance. Neuron 85, 942–958 (2015).

14. Zeisel, A. et al. Brain structure. Cell types in the mouse cortex and hippocampus revealed by single-cell RNA-seq. Science 347, 1138–1142 (2015).

15. Nudelman, A.S. et al. Neuronal activity rapidly induces transcription of the CREB-regulated microRNA-132, in vivo. Hippocampus 20, 492–498 (2010).

16. Aten, S., Hansen, K.F., Hoyt, K.R. & Obrietan, K. The miR-132/212 locus: a complex regulator of neuronal plasticity, gene expression and cognition. RNA Dis 3 (2016).

17. Kepecs, A. & Fishell, G. Interneuron cell types are fit to function. Nature 505, 318–326 (2014).

18. Tasic, B. et al. Adult mouse cortical cell taxonomy revealed by single cell transcriptomics. Nat Neurosci 19, 335–346 (2016).

19. Rudy, B., Fishell, G., Lee, S. & Hjerling-Leffler, J. Three groups of interneurons account for nearly 100% of neocortical GABAergic neurons. Dev Neurobiol 71, 45–61 (2011).

20. Ebert, D.H. & Greenberg, M.E. Activity-dependent neuronal signalling and autism spectrum disorder. Nature 493, 327–337 (2013).

## References

1. Tasic, B. et al. Adult mouse cortical cell taxonomy revealed by single cell transcriptomics. Nat Neurosci 19, 335–346 (2016).

2. Lake, B.B. et al. Neuronal subtypes and diversity revealed by single-nucleus RNA sequencing of the human brain. Science 352, 1586–1590 (2016).

